# Extinction risk in terrestrial vertebrates is associated with niche limits that reflect climatic vulnerability

**DOI:** 10.64898/2026.06.22.733696

**Authors:** Matthew Nagy-Watson, Jeremy T. Kerr

## Abstract

Anthropogenic climate and land-use change are driving an emerging extinction crisis that is expected to intensify in the future. Species’ climatic niche limits shape their sensitivity to these pressures, potentially leading to disproportionate extinction risk among more climatically vulnerable species. We test whether realized climatic niche limits are associated with current and projected extinction risk across >23,000 terrestrial vertebrate species. We assessed the phylogenetic structure of thermal and aridity niche limits and related them to IUCN threat status and simulated future extinctions. We show that realized niche limits are phylogenetically conserved, indicating evolutionary clustering of climatic tolerances. Species with colder upper thermal limits were more likely to be classified as threatened across taxa. Aridity niche limits show weaker and less consistent relationships with current threat status. Simulated extinction scenarios reveal taxon-specific patterns of climatic niche loss compared to random species extinctions. We also show significant reductions in phylogenetic diversity relative to randomized expectations based on simulated species extinctions. We find that extinction risk is systematically associated with species’ climatic niche limits, reflecting evolutionary constraints on environmental tolerance. These results indicate that future extinctions will disproportionately affect climatically vulnerable lineages, with cascading consequences for phylogenetic diversity and ecosystem functioning.

## Introduction

Extant species represent about 1% of the total number of species estimated to have evolved over the last 3.5 billion years (Raup, 1986). Over most of this period, rates of species extinctions were low (De Vos et al., 2015) and balanced by the emergence of new species (Barnosky et al., 2011). Currently, even conservative estimates show that rates of extinction exceed estimated background rates by 100-fold (Ceballos et al., 2015) and are primarily attributed to anthropogenic pressures (Di Marco et al., 2015). Of the 12 threat classifications used by the International Union for the Conservation of Nature (IUCN) to identify the cause of species declines, eight are directly associated with human activities, such as land use conversion, resource exploitation, and pollution (IUCN Red List of Threatened Species, 2022). Additionally, anthropogenic climate change is altering global temperature and precipitation regimes and already contributes to species declines (Miller-Struttmann, 2024; Soroye et al., 2020; Spooner et al., 2018) and the present-day extinction crisis (Urban, 2024).

Over evolutionary time, climatic variability has imposed selective pressures that have shaped phylogenetic diversity (Theodoridis et al., 2020) and species-specific trait adaptations (Rubalcaba et al., 2023) within and among biological communities. Combined with constraints imposed by biotic interactions (Paquette & Hargreaves, 2021), these adaptations help determine species’ distributions and their realized niches (Sunday et al., 2012; Vandermeer, 1972). Realized niches represent the breadth of environmental conditions and biotic interactions under which species can survive and optimize reproduction (Hargreaves et al., 2014; Helaouët & Beaugrand, 2009; Lee-Yaw et al., 2016). Importantly, realized niche limits derived from range boundaries across terrestrial vertebrates are positively associated with species’ physiological tolerances (Watson & Kerr, 2025a). Further, realized niche limits are inextricably linked to species’ morphological and behavioural traits (Buckley et al., 2015; Helaouët & Beaugrand, 2009; Watson & Kerr, 2025b). As climate change alters global temperature and aridity regimes, differences in realized niche limits between species may reflect susceptibility to climate change driven declines and extinction risk (Soroye et al., 2020; Williams et al., 2022).

Historically, anthropogenic land use conversion has been the primary driver determining extinction risk across taxa through impacts of habitat loss, human-wildlife conflicts, and environmental exposure (Allan et al., 2015; T. M. Brooks et al., 2002; Li et al., 2022). Human land use conversion has affected ∼32% of the global land area between 1960-2019 (Winkler et al., 2021), transforming natural habitats into more environmentally homogeneous landscapes (Newbold et al., 2015). The loss of vegetation diversity and habitat complexity associated with land use conversion often reduces the availability of local microclimates, resulting in hotter and drier environmental conditions (Findell et al., 2017) and increases individual organisms’ exposure to broader regional climatic conditions (Kerr et al., 2025). Consequently, land-use conversion may exert stronger negative effects on species occupying habitats near their environmental tolerance limits, where reduced environmental buffering may leave populations more vulnerable to additional stressors. Previous studies have found that population responses to land-use change are influenced by species’ positions within climatic niche space (Williams et al., 2022; Williams & Newbold, 2021). Species adapted to hotter and drier environmental conditions may therefore be less susceptible to anthropogenic land use conversion than species adapted to cooler and wetter habitats.

In addition to land use change, warming conditions and extreme heat events are pushing many species towards and beyond the upper limits of their thermal niches, which can reduce individual organismal fitness (Kingsolver et al., 2013). As species approach their upper thermal niche limits, they are likely to experience increased risk of heat stress and face greater thermoregulatory demands (Buckley et al., 2015; Takamata, 2012). Over time, these energetic demands and physiological stress can reduce body condition, reproductive output, and survival (Andreasson et al., 2018; du Plessis et al., 2012; Van de Ven et al., 2020; Woodroffe et al., 2017). Further, extreme heat events can result in mass population die-offs over relatively short periods of time (Ratnayake et al., 2019). However, species are unlikely to respond uniformly to warming conditions because evolutionary histories and adaptive traits differ among taxa (Foden et al., 2013; Pacifici et al., 2017). Susceptibility to climate change driven warming is therefore likely to be inversely related to species’ warm niche limits.

Climate change is also altering precipitation regimes, leading to changes in regional aridity intensity (Luo et al., 2023). Current predictions anticipate that under climate change, drylands will comprise more than half of the global land surface area by the end of this century (Huang et al., 2016). Increasing aridity can result in pronounced losses of primary productivity, reduced food and water availability, and substantial habitat loss (Golodets et al., 2015). Intensification of global aridity is consequently predicted to result in over 55% of terrestrial vertebrate species to experience habitat loss due to unprecedented aridity conditions (Liu et al., 2023). Additionally, evidence has shown that concurrent changes in aridity and temperature conditions can interact to exacerbate the impacts of climate change (Watson & Kerr, 2025b; Williams et al., 2022). Although evaporative cooling can help organisms maintain optimal body temperatures under increasingly hot conditions, it can also increase the risk of lethal dehydration when replacement water sources are limited (Albright et al., 2017; Sannolo & Carretero, 2019). Species that are adapted to the pressures of dryland conditions are therefore less likely to be negatively impacted by drying conditions compared to species that have evolved under wetter conditions (Conenna et al., 2021; Zylstra et al., 2025). As the global climate continues to change, species with wetter and cooler niche requirements are therefore likely to exhibit greater extinction risks.

It is possible for species to adapt to persistent climate changes (Gardner et al., 2011; Sheridan and Bickford, 2011; Somero, 2010), but current rates of climate change outpace historic rates of niche evolution (Quintero & Wiens, 2013). Many species therefore likely have limited capacity to adapt to emerging conditions sufficiently rapidly to persist in their present locations. In response, many species have shifted their geographical ranges and phenologies to enable them to remain within more tolerable environmental conditions (Chen et al., 2011; Harris et al., 2018; Zaifman et al., 2017). However, differences in species dispersal capacities and environmental connectivity constrain species’ abilities to track suitable environments (Boyles et al., 2011; Pacifici et al., 2017; Thuiller, 2004), imposing greater local and global risks of extinction for some taxa (Calosi et al., 2007; Williams et al., 2021). Climate change-driven extinctions, whether local or global, will ultimately lead to systematic declines in phylogenetic and functional diversity (Matthews et al., 2024), which threatens ecosystem services and stability (Oliver et al., 2015; Scholes, 2016). Identifying species that are most vulnerable to climate-driven extinction is vital for informing conservation action and mitigating climate change driven loss of biodiversity.

Here, we test whether realized niche limits are associated with extinction risk across terrestrial vertebrates. Specifically, we evaluate whether species’ thermal (hot and cold) and aridity (dry and wet) realized niche limits (Watson & Kerr, 2025a) relate to current and future extinction risk. We first test whether realized niche limits are phylogenetically conserved across 4342 amphibian, 7862 bird, 3623 mammal, and 7814 reptile species, given that these limits are associated with species-specific trait adaptations (D’Agostino et al., 2022; Hantak et al., 2021). We then test whether species’ niche limits are associated with extinction risk by comparing them with current International Union for the Conservation of Nature (IUCN) threat status (IUCN, 2022a), predicting that species with cooler and wetter realized niche limits are more likely to be classified as threatened. We predict that land use change would more negatively impact climatically vulnerable species by reducing microclimate availability, increasing exposure to temperature and aridity extremes more broadly, and consequently imperiling species with cooler and wetter realized niche limits to a greater extent.

We further evaluate whether realized niche limits are associated with projected extinction risk. While threat status provides an estimate of current extinction vulnerability, it does not necessarily reflect future extinction trajectories (Simkins et al., 2025). Differences in generation length can strongly influence demographic responses to climate and land-use change and the rate at which extinction risk accumulates (Bird et al., 2020). Since extinction trajectories are shaped by both threat status and generation length, associations between species’ niche limits and projected extinction risk may differ from those inferred from current threat status. To address this, we generate 100 simulations of predicated species extinctions over the next 100 years for all groups of terrestrial vertebrates, accounting for historic IUCN threat status transition rates and generation length estimates (Andermann et al., 2021). We then test whether realized niche limits differ between modelled extinction sets and random extinction scenarios while accounting for current threat status. We predict that species rendered extinct in these simulations would display colder and wetter realize niche limits compared to random species extinctions. Finally, we assess whether these simulated extinctions result in disproportionate losses of phylogenetic diversity (PD) relative to random extinctions, and thereby more greatly impact ecosystem functions and services (Faith, 2008). Overall, this work evaluates whether historical anthropogenic pressures have disproportionately increased extinction risk among more climatically vulnerable species, whether these patterns are likely to persist under future extinction scenarios, and how such losses will impact phylogenetic diversity and ecosystem functioning.

## Results

### Phylogenetic signal in realized niche limits

We tested for the presence of phylogenetic signals in realized thermal and aridity niche limits for 4342 amphibian, 7930 bird, 3623 mammal, and 7814 reptile species. Pagel’s Lambda (λ) values indicated phylogenetic signals are present for all niche limits in each of these taxonomic groups (Table 1). This indicates that closely related species share relatively similar realized thermal and aridity niche limits, though the strength of this relationship varies across classes of terrestrial vertebrates. For all niche limits, we also generated α estimates based on an Ornstein-Uhlenbeck process which assumes traits are constrained by selection towards on adaptive optima. We then used AIC to compare each model of phylogenetic signal across taxonomic groups. Models using Pagel’s λ better explained estimates of phylogenetic signal of realized niche limits except for lower aridity (dry) niche limits in mammals. Generally, these findings demonstrate that realized niche limits across terrestrial vertebrates are phylogenetic conserved, indicating that similarity in realized niche limits increases with phylogenetic proximity.

**Table 1.** Phylogenetic signal measurements for realized thermal and aridity niche limits for terrestrial vertebrates. Phylogenetic signals are based on Pagel’s Lambda (λ) and an Ornstein-Uhlenbeck (α) evolutionary process model. Negative Δ AIC indicates a Brownian motion (λ) process better explains phylogenetic signal in niche limits.

| *Class* | *Niche Limit* | *Pagel's Lambda ($\lambda$)* | *OU ($\alpha$)* | *$\Delta$ AIC (OU)* |
| --- | --- | --- | --- | --- |
| *Amphibia* | Lower Thermal | 0.814 | 0.039 | -1296.2 |
|  | Upper Thermal | 0.854 | 0.029 | -1041.5 |
|  | Lower Aridity | 0.778 | 0.036 | -873.4 |
|  | Upper Aridity | 0.590 | 0.144 | -872.2 |
| *Aves* | Lower Thermal | 0.693 | 0.041 | -164.2 |
|  | Upper Thermal | 0.751 | 0.034 | -202.0 |
|  | Lower Aridity | 0.516 | 0.061 | -147.3 |
|  | Upper Aridity | 0.747 | 0.039 | -213.1 |
| *Mammalia* | Lower Thermal | 0.855 | 1.079 | -28.3 |
|  | Upper Thermal | 0.846 | 1.208 | -20.5 |
|  | Lower Aridity | 0.753 | 1.762 | 42.2 |
|  | Upper Aridity | 0.883 | 1.365 | 62.5 |
| *Reptilia* | Lower Thermal | 0.786 | 0.061 | -1645.3 |
|  | Upper Thermal | 0.866 | 0.055 | -1635.6 |
|  | Lower Aridity | 0.782 | 0.057 | -1473.9 |
|  | Upper Aridity | 0.663 | 0.153 | -1148.3 |

### Niche limits and current extinction risk

Current IUCN threat status was found to consistently be associated with colder upper thermal niche limits across all classes of terrestrial vertebrates (Figures 1 and 2). Additionally, bird, mammal, and reptile species with warmer lower thermal niche limits were more likely to be classified as threatened (Table 2), with no association observed for amphibians (OR = 0.921, 95% CI = 0.811 – 1.047, p = 0.208). Fewer significant associations between aridity niche limits and threat status were observed compared to thermal niche limits (Figure 1). Contrary to predictions, probability of being classed as threatened decreased with wetter lower aridity (dry) niche limits for amphibians (OR = 0.901, 95% CI = 0.829 – 0.981, p = 0.016) and birds (OR = 0.803, 95% CI = 0.738 – 0.873, p < 0.001), while no association was found for mammals or reptiles (Table 2). Species upper aridity (wet) niche limits were not significantly associated with threat status for any terrestrial vertebrate group. Finally, all four taxonomic groups showed lower probability of being classified as threatened as species range area increased (Figure 1), though this was expected based on the IUCN methods for assigning threat status (IUCN, 2022b).

**Figure 1.**
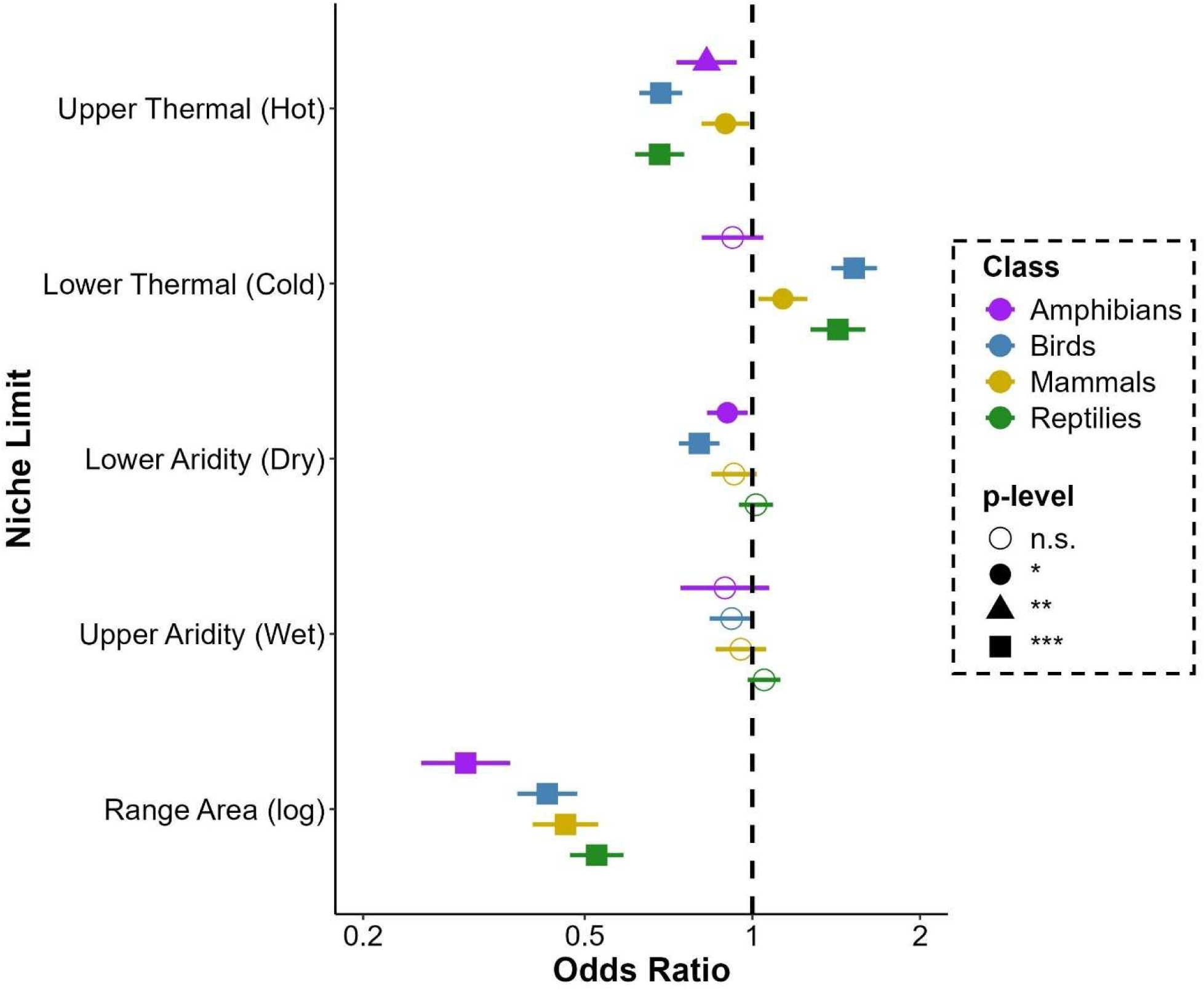
Results of phylogenetic logistic regression comparing threat status to niche limits. Displays the odds ratio for all niche limit variables and log_10_ range area for each taxonomic class. Values less than 1 indicate an increase in the variable lowers probability of being classed as “Threatened”, while values above 1 indicate an increased probability. Figure represents the logistic regression results presented in Table 2. Significance indicated by values not overlapping 0 (solid fill), shape indicates p-level.

**Figure 2.**
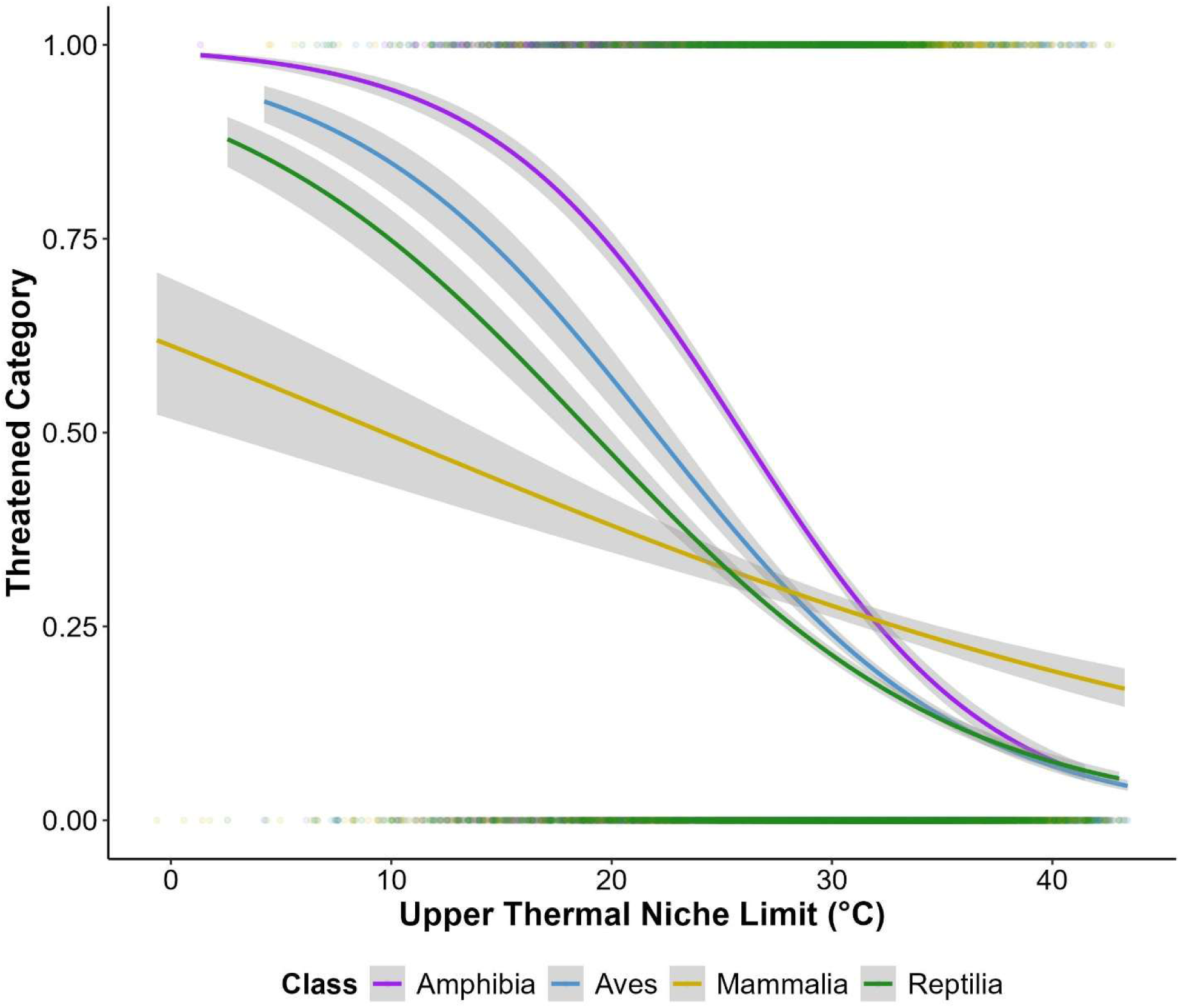
Logistic regression plot of threat status based on upper thermal niche limit. Displays the relationship between change in upper thermal niche limit and probability of being classified as threatened (1) or not threatened (0) for each taxonomic class. Probability of being classified as threatened significantly declines as upper thermal niche limits increases for: Amphibians (β = -0.189, p = 0.003), Birds (β = -0.379, p < 0.001), Mammals (β = -0.111, p = 0.028), and Reptiles (β = -0.383, p < 0.001).

**Table 2.** Phylogenetic logistic regression results evaluating the association between environmental niche limits and IUCN threatened status. All standardized estimates represent the impact of a 1SD change in the respective variable. Likelihood R^2^ reported for each model.

| Amphibia: $R^2_{lik} = 0.465$ | | | | |
| --- | --- | --- | --- | --- |
|  | Estimate | SE | Z | p |
| Intercept | -2.112 | 0.211 | -10.03 | < 0.001 |
| T<sub>max</sub> | -0.189 | 0.064 | -2.98 | 0.003 |
| T<sub>min</sub> | -0.082 | 0.065 | -1.26 | 0.208 |
| A<sub>min</sub> | -0.104 | 0.043 | -2.41 | 0.016 |
| A<sub>max</sub> | -0.114 | 0.094 | -1.22 | 0.224 |
| Log<sub>10</sub> Range Area | -1.186 | 0.094 | -12.61 | < 0.001 |
| Aves: $R^2_{lik} = 0.379$ | | | | |
|  | Estimate | SE | Z | p |
| Intercept | -1.588 | 0.146 | -10.84 | < 0.001 |
| T<sub>max</sub> | -0.379 | 0.045 | -8.35 | < 0.001 |
| T<sub>min</sub> | 0.421 | 0.048 | 8.70 | < 0.001 |
| A<sub>min</sub> | -0.220 | 0.043 | -5.14 | < 0.001 |
| A<sub>max</sub> | -0.086 | 0.046 | -1.86 | 0.062 |
| Log<sub>10</sub> Range Area | -0.848 | 0.063 | -13.41 | < 0.001 |
| Mammalia: $R^2_{lik} = 0.455$ | | | | |
|  | Estimate | SE | Z | p |
| Intercept | -0.634 | 0.239 | -2.65 | 0.008 |
| T<sub>max</sub> | -0.111 | 0.051 | -2.20 | 0.028 |
| T<sub>min</sub> | 0.126 | 0.052 | 2.42 | 0.015 |
| A<sub>min</sub> | -0.076 | 0.048 | -1.59 | 0.111 |
| A<sub>max</sub> | -0.048 | 0.053 | -0.90 | 0.370 |
| Log<sub>10</sub> Range Area | -0.773 | 0.069 | -11.16 | < 0.001 |
| Reptilia: $R^2_{lik} = 0.286$ | | | | |
|  | Estimate | SE | Z | p |
| Intercept | -1.656 | 0.067 | -24.73 | < 0.001 |
| T<sub>max</sub> | -0.383 | 0.052 | -7.39 | < 0.001 |
| T<sub>min</sub> | 0.353 | 0.058 | 6.09 | < 0.001 |
| A<sub>min</sub> | 0.015 | 0.036 | 0.41 | 0.680 |
| A<sub>max</sub> | 0.048 | 0.035 | 1.40 | 0.161 |
| Log<sub>10</sub> Range Area | -0.644 | 0.056 | -11.44 | < 0.001 |

### Future extinction scenario comparison

Thermal niche limits differed significantly between extinction simulations based on anticipated ICUN status transitions and generation lengths compared to threat status-controlled random extinctions (Tables 3 and 4). Under the IUCN scenario, species had overall colder upper thermal niche limits for birds (M_EMM_ = -0.298, 95% CI = -0.359 – - 0.238) and mammals (M_EMM_ = -0.341, 95% CI = -0.410 – -0.272) compared to the random scenario (Figure 3; Table 7). However, hotter upper thermal niche limits were found for amphibian (M_EMM_ = 0.146, 95% CI = 0.113 – 0.178) and reptile (M_EMM_ = 0.716, 95% CI = 0.670 – 0.762) extinctions under the IUCN scenario compared to the random scenario (Figure 3; Table 7). In contrast, hotter lower thermal niche limits were observed for birds (M_EMM_ = 0.855, 95% CI = 0.754 – 0.955), mammals (M_EMM_ = 1.153, 95% CI = 1.039 – 1.267), and reptiles (M_EMM_ = 0.732, 95% CI = 0.655 – 0.808) under the IUCN scenario, while amphibians exhibited colder lower thermal niche limits (M_EMM_ = -0.235, 95% CI = - 0.288 – -0.181) (Figure 3; Table 7).

**Figure 3.**
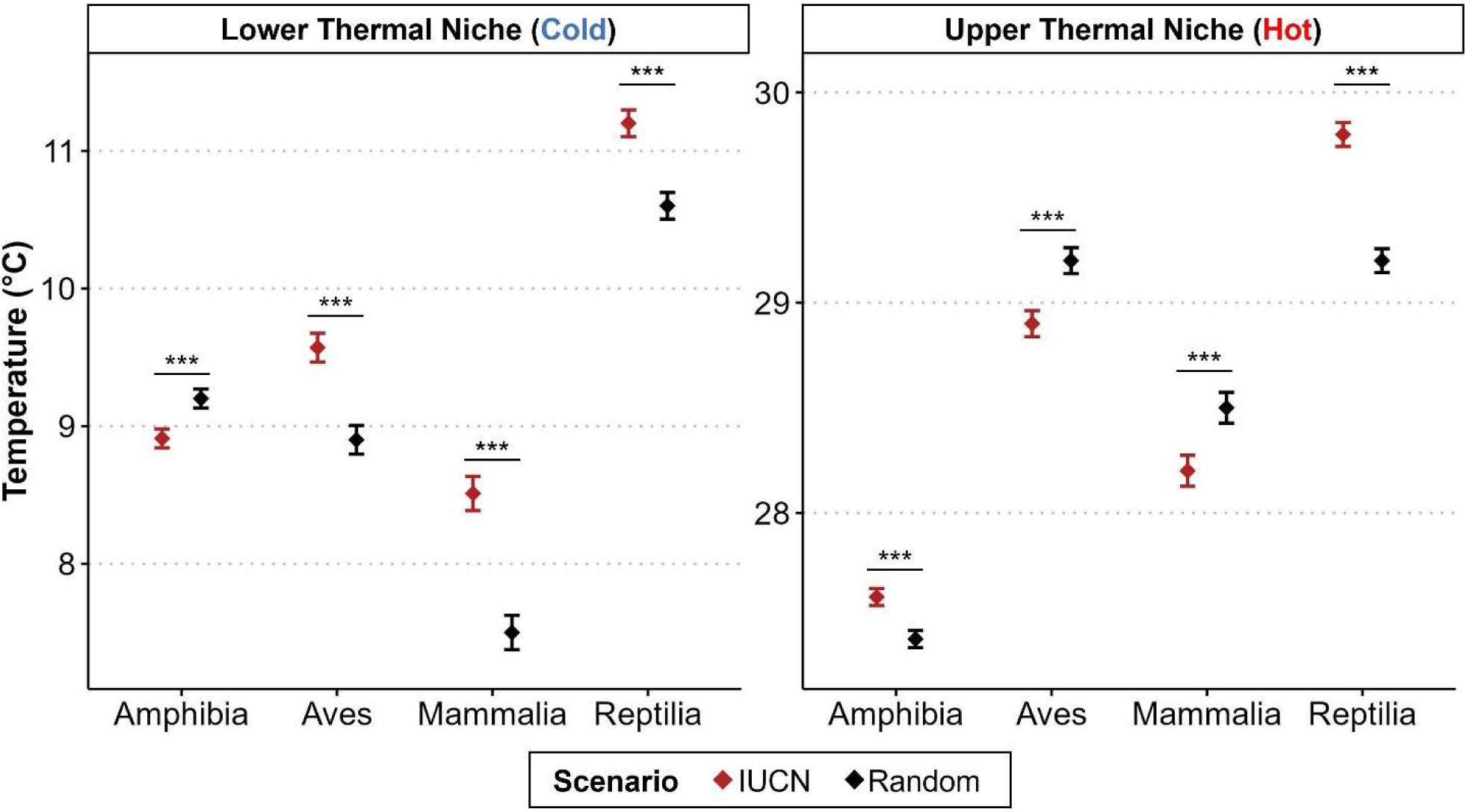
Linear mixed effect model results comparing thermal niche limits between extinction scenarios. Red values indicate standardized temperature values for extant species after the removal of extinctions generated by iucnsim, while black values represent standardized temperature values for extant species after the random removal of species. Results are representative of results presented in Table 4.3 & 4.4. The number of species extinctions are consistent across threat status for both IUCN and random scenarios. Asterisk indicates significant difference in temperature between extinction scenarios (p-level: * 0.01, ** 0.001, *** 0.001).

**Table 3.** Linear mixed effect model results comparing upper thermal (hot) niche limits between extinction scenarios. Marginal/Conditional R^2^ = 0.267/0.275.

| Fixed Effects |  |  |  |  |
| --- | --- | --- | --- | --- |
|  | Estimate | Std. Error | z value | p value |
| Intercept (Class: Amphibia; Scenario: IUCN) | 22.162 | 0.317 | 70.00 | < 0.001 |
| Scenario: Random | -0.190 | 0.020 | -9.34 | < 0.001 |
| Class: Aves | 1.294 | 0.027 | 47.76 | < 0.001 |
| Class: Mammalia | 0.592 | 0.031 | 19.37 | < 0.001 |
| Class: Reptilia | 2.179 | 0.024 | 89.20 | < 0.001 |
| Log Range Area | 1.543 | 0.005 | 292.53 | < 0.001 |
| Scenario: Random x Class: Aves | 0.523 | 0.037 | 14.26 | < 0.001 |
| Scenario: Random x Class: Mammalia | 0.531 | 0.042 | 12.73 | < 0.001 |
| Scenario: Random x Class: Reptilia | -0.457 | 0.035 | -13.22 | < 0.001 |
| Random Effects |  |  |  |  |
|  | Group | Variance | SD | N obs. |
| Sample Number | Intercept | < 0.0001 | 0.0002 | 100 |
| IUCN Category | Intercept | 0.1991 | 0.4463 | 2 |
| Residual |  | 18.9700 | 4.3557 | 422013 |

**Table 4.** Linear mixed effect model results comparing lower thermal (cold) niche limits between extinction scenarios. Marginal/Conditional R^2^ = 0.274/0.274.

| Fixed Effects |  |  |  |  |
| --- | --- | --- | --- | --- |
|  | Estimate | Std. Error | z value | p value |
| Intercept (Class: Amphibia; Scenario: IUCN) | 18.544 | 0.117 | 158.73 | < 0.001 |
| Scenario: Random | 0.256 | 0.034 | 7.43 | < 0.001 |
| Class: Aves | 0.661 | 0.046 | 14.36 | < 0.001 |
| Class: Mammalia | -0.399 | 0.052 | -7.68 | < 0.001 |
| Class: Reptilia | 2.238 | 0.041 | 53.95 | < 0.001 |
| Log Range Area | -2.737 | 0.009 | -305.37 | < 0.001 |
| Scenario: Random x Class: Aves | -0.933 | 0.062 | -14.99 | < 0.001 |
| Scenario: Random x Class: Mammalia | -1.255 | 0.071 | -17.73 | < 0.001 |
| Scenario: Random x Class: Reptilia | -0.847 | 0.059 | -14.43 | < 0.001 |
| Random Effects |  |  |  |  |
|  | Group | Variance | SD | N obs. |
| Sample Number | Intercept | < 0.0001 | < 0.0001 | 100 |
| IUCN Category | Intercept | 0.0246 | 0.1568 | 2 |
| Residual |  | 54.6900 | 7.3950 | 422013 |

Differences in aridity niche limits between extinction scenarios showed variable patterns across taxonomic groups (Tables 5 and 6). Compared to the random extinction scenario, amphibians had wetter lower aridity (dry) niche limits (M_EMM_ = 0.012, 95% CI = 0.005 – 0.014) and dryer upper aridity (wet) niche limits (M_EMM_ = -0.104, 95% CI = - 0.127 – -0.082) under the IUCN scenario (Figure 4, Table 7). Both bird and mammal species showed no difference in lower aridity (dry) niche limits between extinction scenarios (Figure 4; Table 7), though both displayed dryer upper aridity (wet) niche limits under the IUCN scenario (bird: M_EMM_ = -0.370, 95% CI = -0.365 – -0.280; mammal: M_EMM_ = -0.506, 95% CI = -0.523 – -0.426). Finally, no significant difference in lower or upper aridity niche limits were found between extinction scenarios for reptiles (Figure 4; Table 7). All analyses included range area as a covariate to control for potential spurious associations between niche limits and extinction scenarios, since range area informs IUCN threat status and can impact niche limit breadth.

**Figure 4.**
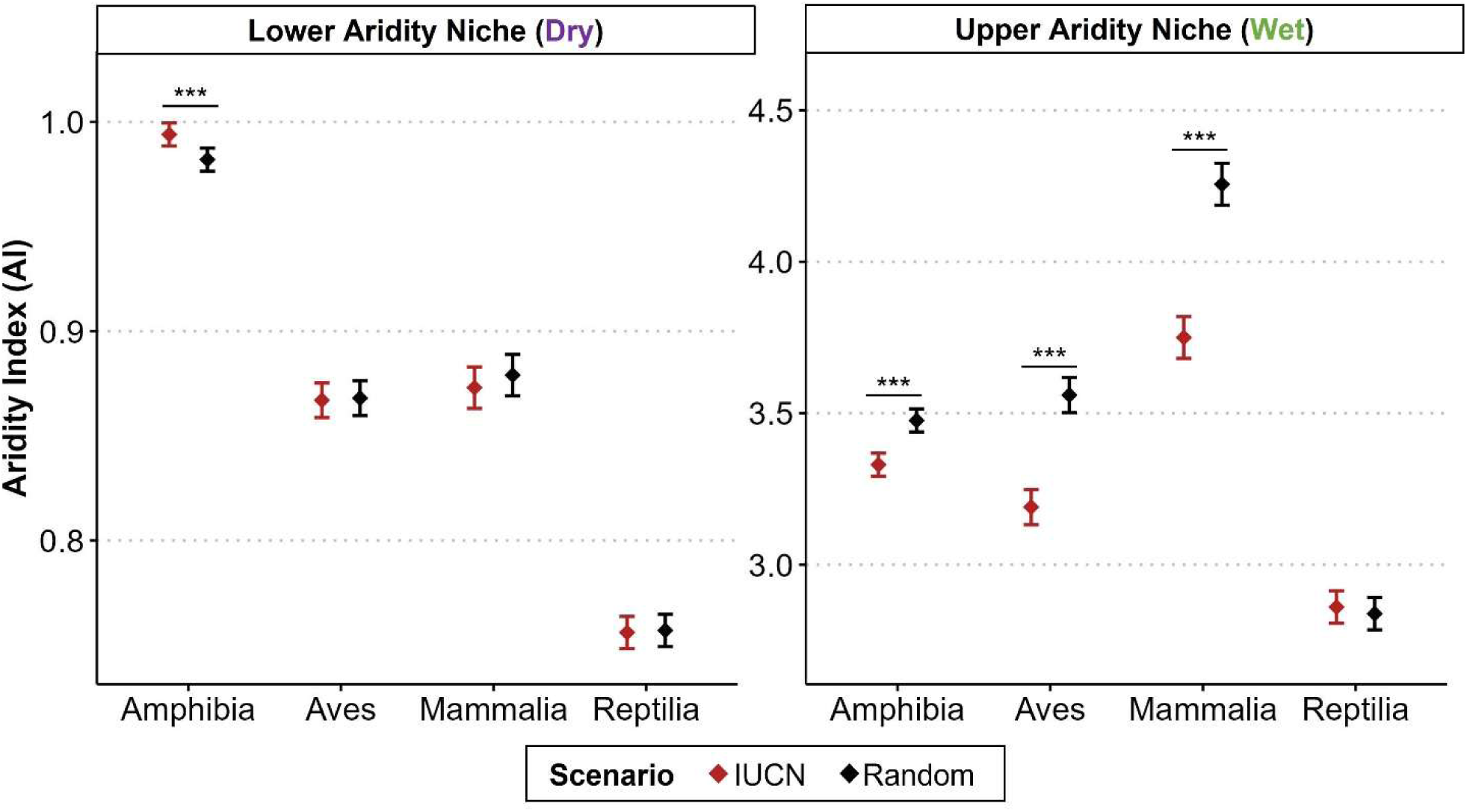
Linear mixed effect model results comparing aridity niche limits between extinction scenarios. Red values indicate standardized Aridity Index values for extant species after the removal of extinctions generated by iucnsim, while black values represent standardized Aridity Index values for extant species after the random removal of species. Results are representative of results presented in Table 4.5 & 4.6. The number of species extinctions are consistent across threat status for both IUCN and random scenarios. Asterisk indicates significant difference in Aridity Index between extinction scenarios (p-level: * 0.01, ** 0.001, *** 0.001).

**Table 5.** Linear mixed effect model results comparing upper aridity (wet) niche limits between extinction scenarios. Marginal/Conditional R^2^ = 0.114/0.114.

| Fixed Effects |  |  |  |  |
| --- | --- | --- | --- | --- |
|  | Estimate | Std. Error | z value | p value |
| Intercept (Class: Amphibia; Scenario: IUCN) | 0.328 | 0.029 | 11.25 | < 0.001 |
| Scenario: Random | 0.146 | 0.019 | 7.64 | < 0.001 |
| Class: Aves | -0.132 | 0.025 | -5.18 | < 0.001 |
| Class: Mammalia | 0.427 | 0.029 | 14.86 | < 0.001 |
| Class: Reptilia | -0.461 | 0.023 | -20.06 | < 0.001 |
| Log Range Area | 0.852 | 0.005 | 170.49 | < 0.001 |
| Scenario: Random x Class: Aves | 0.225 | 0.034 | 6.52 | < 0.001 |
| Scenario: Random x Class: Mammalia | 0.360 | 0.039 | 9.19 | < 0.001 |
| Scenario: Random x Class: Reptilia | -0.168 | 0.032 | -5.17 | < 0.001 |
| Random Effects |  |  |  |  |
|  | Group | Variance | SD | N obs. |
| Sample Number | Intercept | < 0.0001 | 0.0002 | 100 |
| IUCN Category | Intercept | 0.0009 | 0.0292 | 2 |
| Residual |  | 16.7600 | 4.0934 | 421994 |

**Table 6.**
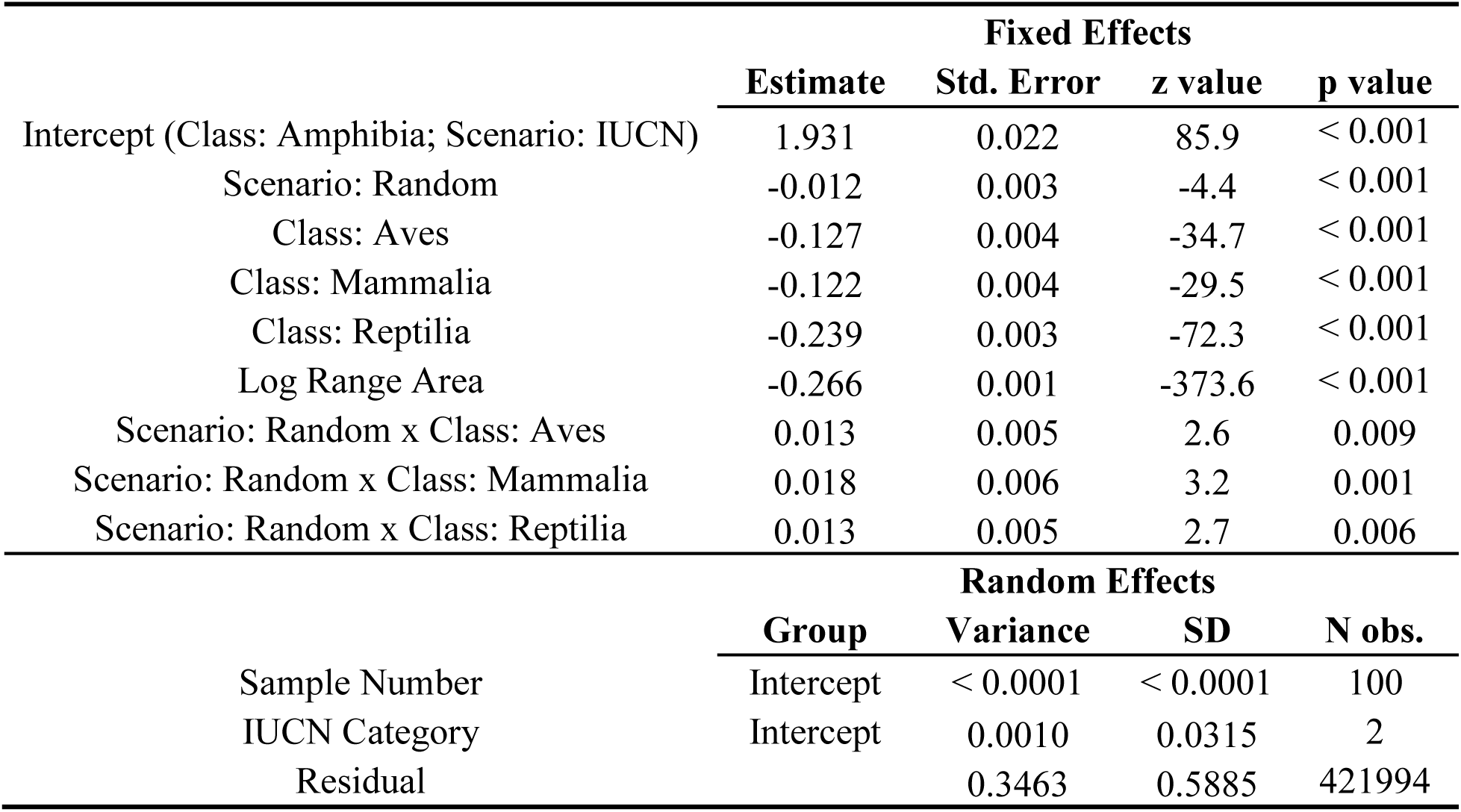
Linear mixed effect model results comparing lower aridity (dry) niche limits between extinction scenarios. Marginal/Conditional R^2^ = 0.349/0.351.

Niche limits were found to exhibit significant phylogenetic signals in my previous analyses (Table 1) and were associated with extinction risk (Tables 2-7). This indicates the potential for phylogenetically clustered species extinctions under my IUCN scenario, which could lead to elevated losses of PD. Observed levels of extant PD differed significantly between IUCN and random extinction scenarios for all taxonomic groups (Table 8, Figure 5). Lower levels of extant PD were observed for amphibians (M_EMM_ = - 1.254, 95% CI = -1.512 – -0.995), mammals (M_EMM_ = -0.315, 95% CI = -0.574 – - 0.0572), and reptiles (M_EMM_ = -0.746, 95% CI = -1.004 – -0.4874) under the IUCN extinction scenarios compared to random extinctions. However, remaining extant PD for birds was significantly greater under IUCN extinction scenarios compared to random scenarios (M_EMM_ = 0.775, 95% CI = 0.516 – 1.033).

**Figure 5.**
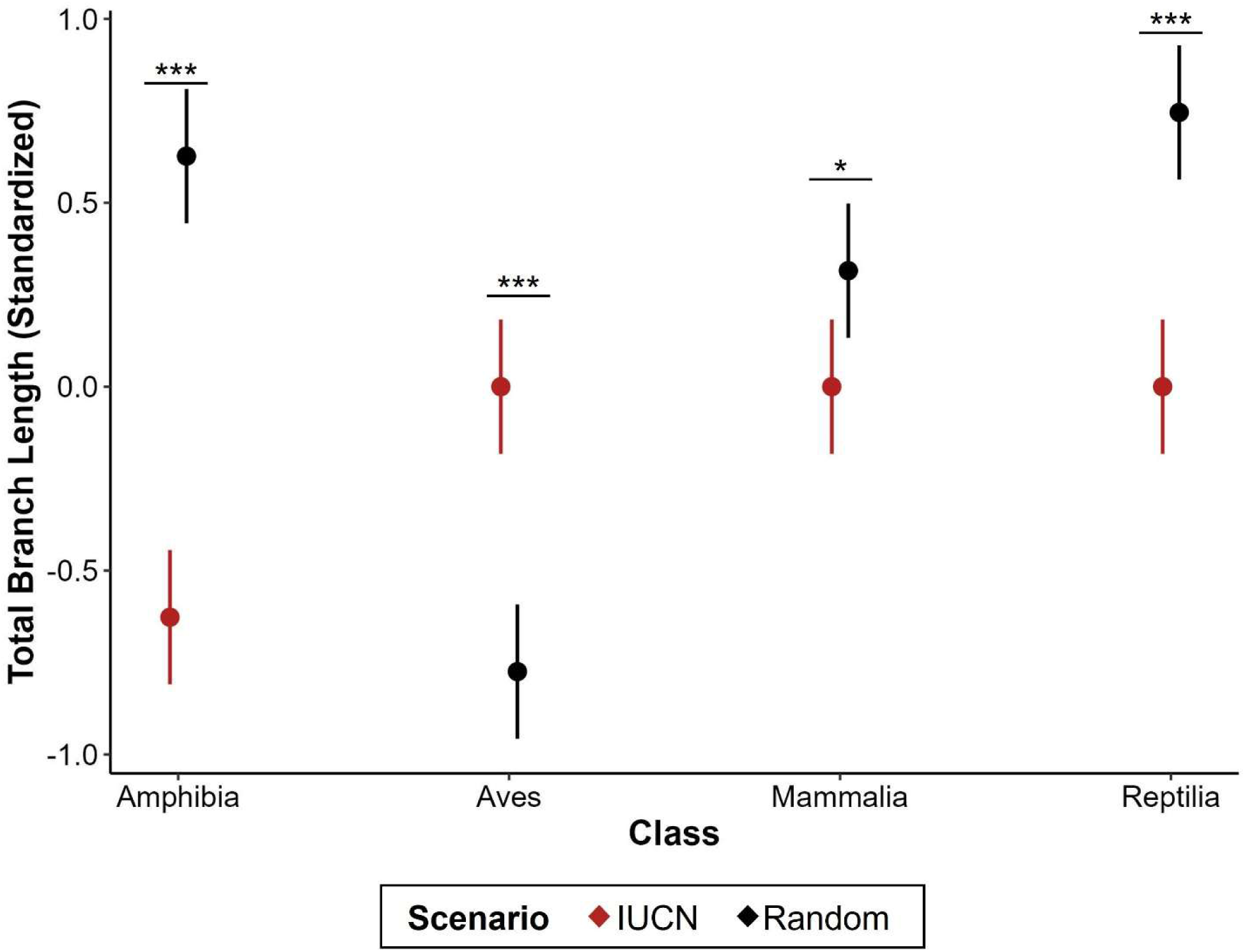
Linear model results comparing phylogenetic diversity between extinction scenarios. PD was calculated as sum total branch length of extant species. Red values indicate standardized PD for extant species after the removal of extinctions generated by *iucnsim*, while black values represent standardized PD for extant species after the random removal of species. The number of species extinctions are consistent across threat status for both IUCN and random scenarios. Asterisk indicates significant difference in PD between extinction scenarios (p-level: * 0.01, ** 0.001, *** 0.001).

**Table 7.** Estimated marginal means of niche limit contrasts between extinction scenarios. Estimates represent the marginal means of IUCN niche limit – random niche limit, with positive estimates indicating hotter/wetter niche limits for IUCN extinct species. Contrasts were generated based on linear mixed effect model results presented in Tables 4-7.

| Class | Niche | Estimate | SE | df | T value | p value |
| --- | --- | --- | --- | --- | --- | --- |
| *Amphibia* | T<sub>min</sub> | -0.235 | 0.034 | 422002 | -7.43 | < 0.001 |
|  | T<sub>max</sub> | 0.146 | 0.020 | 422002 | 9.34 | < 0.001 |
|  | A<sub>min</sub> | 0.012 | 0.003 | 421983 | 4.38 | < 0.001 |
|  | A<sub>max</sub> | -0.104 | 0.019 | 421983 | -7.64 | < 0.001 |
| *Aves* | T<sub>min</sub> | 0.855 | 0.052 | 422002 | 12.94 | < 0.001 |
|  | T<sub>max</sub> | -0.298 | 0.031 | 422002 | -10.81 | < 0.001 |
|  | A<sub>min</sub> | -0.001 | 0.004 | 421983 | -0.23 | 0.819 |
|  | A<sub>max</sub> | -0.323 | 0.029 | 421983 | -12.78 | < 0.001 |
| *Mammalia* | T<sub>min</sub> | 1.153 | 0.062 | 422002 | 16.04 | < 0.001 |
|  | T<sub>max</sub> | -0.341 | 0.037 | 422002 | -9.30 | < 0.001 |
|  | A<sub>min</sub> | -0.006 | 0.005 | 421983 | -1.19 | 0.234 |
|  | A<sub>max</sub> | -0.475 | 0.035 | 421983 | -14.66 | < 0.001 |
| *Reptilia* | T<sub>min</sub> | 0.732 | 0.048 | 422002 | 12.24 | < 0.001 |
|  | T<sub>max</sub> | 0.716 | 0.028 | 422002 | 22.73 | < 0.001 |
|  | A<sub>min</sub> | -0.001 | 0.004 | 421983 | -0.20 | 0.840 |
|  | A<sub>max</sub> | 0.074 | 0.027 | 421983 | 0.84 | 0.403 |

**Table 8.** Linear regression results comparing phylogenetic diversity between extinction scenarios and taxonomic classes. Multiple/Adjusted R^2^ = 0.228/0.222. Residual standard error = 0.930 on 792 degrees of freedom. Contrasts represent the estimated marginal mean difference of PD between IUCN and random extinctions for each taxonomic class. No collinearity between variables was observed based on VIF < 5: Scenario = 2.00, Class = 1.41, Scenario x Class = 1.65.

|  | Estimate | Std. Error | z value | p value |  |
| --- | --- | --- | --- | --- | --- |
| Intercept (Class: Amphibia; Scenario: IUCN) | -0.627 | 0.093 | -6.74 | < 0.001 |  |
| Scenario: Random | 1.254 | 0.132 | 9.53 | < 0.001 |  |
| Class: Aves | 0.627 | 0.132 | 4.76 | < 0.001 |  |
| Class: Mammalia | 0.627 | 0.132 | 4.76 | < 0.001 |  |
| Class: Reptilia | 0.627 | 0.132 | 4.76 | < 0.001 |  |
| Scenario: Random x Class: Aves | -2.028 | 0.186 | -10.90 | < 0.001 |  |
| Scenario: Random x Class: Mammalia | -0.938 | 0.186 | -5.04 | < 0.001 |  |
| Scenario: Random x Class: Reptilia | -0.508 | 0.186 | -2.73 | 0.006 |  |
| Contrast: IUCN - Random |  |  |  |  |  |
| Class | Estimate | SE | df | T value | p value |
| Amphibia | -1.254 | 0.132 | 792 | -9.53 | < 0.001 |
| Aves | 0.775 | 0.132 | 792 | 5.89 | < 0.001 |
| Mammalia | -0.315 | 0.132 | 792 | -2.40 | 0.017 |
| Reptilia | -0.746 | 0.132 | 792 | -5.67 | < 0.001 |

## Discussion

Our results demonstrate that realized niche limits are consistently associated with present day extinction risk as assessed by the IUCN. We show that, after controlling for range size, species with narrower and cooler thermal niches face greater threats at present than species with broader niches and higher upper thermal limits. To date, population declines and threat-status up listing (e.g., reclassification of a near threatened species as vulnerable) of species have been attributed largely to habitat loss and overexploitation caused by human activities (Hogue & Breon, 2022), while climate change is an emerging contributor (Luedtke et al., 2023; Woo-Durand et al., 2020). However, land use change can expose organisms to both higher daytime maximum and lower nighttime minimum temperatures compared to those experienced in intact habitats (De Frenne et al., 2019), which compounds impacts of habitat loss and climate change-driven warming (Findell et al., 2017). Our reported associations between thermal niche limits and extinction risk indicate that climate change is likely a meaningful, and perhaps even underestimated, contributor to recent species declines.

Across vertebrate taxa, species with colder upper thermal niche limits were more likely to be classified as threatened, highlighting that extinction vulnerability is currently associated with species’ thermal niche. Further, species with warmer lower thermal niche limits showed higher extinction risk in birds, mammals, and reptiles, suggesting that thermal breadth and position within thermal space may jointly structure extinction vulnerability. Amphibians, however, showed no association with lower thermal niche limits, indicating that their extinction risk may be driven more strongly by other environmental constraints or life-history factors. Likely, species with warmer and broader thermal niche limits have historically been less negatively impacted by the thermal conditions associated with land use and climate change, driving observed associations between thermal niche limits and current extinction risk. Overall, these patterns align with predictions that more thermally vulnerable species are disproportionately represented among currently threatened taxa.

Present day associations between thermal niche limits and threat status results in greater projected extinction risk for thermally vulnerable bird and mammal species. Under IUCN-based extinction simulations extinct bird and mammal species show cooler upper thermal niche limits compared to randomly selected extinctions, which is consistent with current threat patterns. In contrast, amphibians and reptiles show the opposite pattern, with extinct species exhibiting warmer upper thermal niche limits under IUCN simulations than under random extinction scenarios. These contrasting patterns likely reflect differences in the current distribution of extinction risk, dispersal capacity, and physiology among endothermic (birds and mammals) and ectothermic (amphibians and reptiles) vertebrate groups (Hillman et al., 2014). Birds and mammals generally exhibit greater mobility and broader dispersal capacities (Hancock & Hedrick, 2018), allowing them to track shifting climatic and land use conditions more effectively. Consequently, extinction risk in these groups is associated more broadly with factors beyond temperature, including annual primary productivity, body size, range size, and human population density (Davidson et al., 2017; Munstermann et al., 2022a).

In contrast, amphibians and many reptiles have more limited dispersal capacity and greater physiological dependence on microclimatic conditions (Taylor et al., 2021). These constraints are particularly pronounced in tropical lowland systems, where species are already in closer proximity to their niche limits, have limited capacity to escape warming or drying trends, and are facing intense land use pressures (Alroy, 2015; Frishkoff et al., 2015; Mi et al., 2023; Pottier et al., 2025). Together with restricted geographic ranges and short generation lengths (Mancini et al., 2025; Mi et al., 2023), these factors may explain the contrasting patterns in thermal niche limit differences between extinction scenarios (IUCN-based versus random) observed in amphibians and reptiles relative to birds and mammals. These results indicate patterns of association between niche limits and extinction risk may diverge relative to present day extinction risk for ectothermic vertebrates, while remaining consistent for endothermic species.

Associations between aridity niche limits and present-day extinction risks are weaker and less consistent compared to thermal niche limits. I only detected associations with current threat status and dry niche limits in birds and amphibians, and these were contrary to initial predictions. Wetter lower aridity niche limits were associated with reduced threat status probability for these groups, while upper aridity limits showed no relationship with extinction risk across all taxa. For amphibians and birds, these results demonstrate that species characterized by niche limits at the drier end of the aridity index gradient are more likely to be classified as threatened.

The association between dryer aridity niche limits and increased probability of being classified as threatened could be related to the effects of land-use conversion, which can be more severe drylands (García-Vega & Newbold, 2019). The limited water availability in drylands, combined with reduced vegetation cover and lower microclimate stability associated with land-use change can increase species sensitivity to anthropogenic disturbance and exposure to prevailing climatic conditions (Morant et al., 2025). This would result in observed patterns of aridity niche limit associations with extinction risk in amphibians and birds, though it does not explain the lack of association in mammals and reptiles. In dryland environments, amphibians and birds have been found to be more exposed to prevailing climatic conditions (Riddell et al., 2021; Zhang et al., 2021) and have greater water demands (Taylor et al., 2021; Wu et al., 2024), when compared to mammals and reptiles. Birds and amphibians occupying dryland environments may therefore be disproportionately vulnerable to climate and land-use pressures, providing a potential explanation for the association between lower aridity niche limits and threat status observed in these taxa compared with no association in mammals and reptiles. Overall, aridity niche limits predict present-day threat status poorly, indicating thermal vulnerability is more informative when evaluating extinction risk.

Aridity niche limits also show weaker and more heterogeneous effects across extinction simulations compared to thermal niche limits. Reptile aridity niche limits show no significant difference between extinction scenarios (IUCN-based and random), which is concordant with patterns observed for present-day extinction risk. However, under IUCN extinction scenarios amphibians, birds, and mammals show overall narrower aridity niche limits compared to random extinctions. Specifically, amphibians with wetter minimum aridity requirements were more likely to go extinct within simulated extinctions, indicating that the least dry tolerant species are at greatest risk of extinction. Since amphibian life histories are more reliant on water availability compared to the other vertebrate groups (Shoemaker, 1988), it tracks that species with lower aridity tolerance would be most vulnerable to historic land use and climate change pressures (Nowakowski et al., 2017). This would then result in these less arid tolerant species being more likely to trend towards extinction under my simulations. Additionally, we also observed significantly dryer wet niche limits (upper aridity) in amphibians, birds, and mammals, in contrast to no observed associations in present day extinction risk. Likely, species within these three taxonomic groups that occupy drier overall environments are more vulnerable to current land use pressures, as highlighted previously, which would result in increased risk of extinction within my simulations. Further, species that occupy drier environments tend to display smaller overall body size (Gouveia & Correia, 2016; Watson & Kerr, 2025b), which is associated with shorter generation lengths (Cejp & Griebeler, 2024; Gaillard et al., 2005). Combined, the vulnerability to land use conversion in dryland species and shorter generation lengths potentially put species with dryer overall niches at greater risk of extinction within my simulations.

Realized thermal and aridity niche limits are phylogenetically structured across terrestrial vertebrates. In other words, climatic tolerances (as inferred from realized niche observations) show phylogenetic niche conservatism and vulnerability to climate change is evolutionarily clustered, rather than randomly distributed across the phylogenies of vertebrate taxa. In general, extinction risk has a significant phylogenetic signal across terrestrial vertebrates (Corey & Waite, 2008; Yessoufou et al., 2012), highlighting that vulnerability is potentially associated with phylogenetically structured trait adaptations (Greenberg et al., 2021; Munstermann et al., 2022b). Species with similar morphological traits will exhibit more similar niche limits (Barnagaud et al., 2014; Petit et al., 2025) and by extension, sensitivity to climate and land use changes, which is also likely to cause extinction risk to be clustered across taxa. Consequently, the link between of extinction risk and realized niche limits could negatively impact the most climatically vulnerable species, which is likely to further elevate their extinction risks under continued climate change.

Since niche limits with respect to temperature and aridity are phylogenetically conserved and associated with extinction risk, future extinctions should also be phylogenetically clustered. Clustered extinctions can cause disproportionate losses of phylogenetic diversity loss due to the elimination of evolutionarily distinct branches from a taxon’s phylogenetic tree (Ali et al., 2023; Matthews et al., 2024). Our findings support that view, with PD losses differing significantly between IUCN simulations and randomized extinction scenarios for all taxa. However, the direction of PD change is not uniform. We observe reduced extant PD under IUCN-based extinctions compared to random in amphibians, mammals, and reptiles. This indicates that simulated extinctions based on IUCN threat status transitions and species-specific generation lengths result in greater negative impacts on diversity loss relative to random species extinctions. In contrast, birds show greater remaining extant PD under IUCN extinction simulations than under a random extinction scenario. One potential explanation for this is that bird species at greatest risk are less phylogenetically clustered leading to fewer instances of loss of distinct evolutionary lineages at higher taxonomic levels (Genus, Family, etc.) (Parhar & Mooers, 2011). It is also possible phylogenetically clustered extinctions are occurring in lineages of birds with a large number of species, and that too few species go extinct over the 100 years specified in my simulations to result in the elimination of higher order groups. Clustered extinctions in closely related species with no loss of higher order taxonomic groupings would explain why PD loss was less severe under IUCN simulations compared to random extinctions observed in birds (Parhar & Mooers, 2011). Given a longer time period for extinctions we would then expect patterns of PD loss for birds to converge with those observed for other taxa.

Patterns of PD loss between extinction scenarios (IUCN based and random) vary across vertebrate groups. However, species extinctions will invariably result in declining PD, which has important ecological consequences. More phylogenetically diverse assemblages are typically associated with greater functional diversity (Tucker et al., 2018), meaning that non-random extinction of climatically vulnerable lineages is likely to have cascading effects on ecosystem functioning (Oliver et al., 2015). While we did not directly model functional diversity, the observed structure of PD loss suggests that future extinctions will translate to significant declines in functional diversity. In particular, concentrated losses in amphibians, mammals, and reptiles under IUCN scenarios imply disproportionate erosion of evolutionary history in these groups, with potential consequences for ecosystem functioning and resilience.

We evaluate whether extinction risk inferred from present-day threat and demographic processes aligns with climatic niche structure using time-averaged projections based on IUCN status transition probabilities and species-specific generation lengths (Andermann et al., 2021). It is important to note that our simulations do not represent true extinction rates, nor do they explicitly account for climate or land use change trajectories. Rather, we show that under climate- and land use-agnostic scenarios, extinction risk has significant potential to negatively impact the most climatically vulnerable species. Our findings are consistent with the idea that climatic niche limits are linked to evolutionary history, ecological tolerance, and extinction vulnerability. Previous studies have shown that species near their thermal and hydric limits experience declines in occupancy, abundance, and fitness as climate changed (Soroye et al., 2020; Williams et al., 2022; Williams & Newbold, 2021). Our results extend this conclusion by showing that such niche constraints are not only associated with current extinction risk but also shape projected extinction pathways under simulated extinction scenarios.

Globally, species are already experiencing declines in abundance and local population extinctions associated with recent climate change and land use pressures (Ferrante et al., 2025; Newbold et al., 2025; Soroye et al., 2020; Spooner et al., 2018). Our results suggest that these pressures will continue to disproportionately affect vertebrate species that are less heat tolerant and display narrower thermal and aridity niche limits. As climate change intensifies, this will likely reinforce non-random extinction patterns across terrestrial vertebrates and drive systematic losses of phylogenetic diversity (Matthews et al., 2024), with implications for ecosystem functioning and services (Oliver et al., 2015; Pecl et al., 2017). Finally, realized niche limits provide help predict extinction risk across taxa. They capture long-term evolutionary constraints on environmental tolerance and vulnerability, making them a potentially valuable tool for identifying species at greatest risk under ongoing environmental change. Incorporating such metrics into conservation planning could improve prioritization by identifying lineages and regions where climatic vulnerability and extinction risk are most strongly aligned.

## Methods

### Realized niche limit and extinction risk data

We obtained yearly mean thermal and aridity realized niche limits for 7,958 amphibian, 11,044 bird, 5,655 mammal, and 9,284 reptile species (Watson and Kerr, 2025). For each species, we also extracted total geographic range area (km²) using expert-informed range maps (BirdLife International and Handbook of the Birds of the World, 2021; IUCN, 2022a). To account for shared evolutionary history in subsequent analyses, we obtained phylogenies for amphibians, birds, mammals, and reptiles from published sources (Jetz et al., 2012; Jetz & Pyron, 2018; Tonini et al., 2016; Upham et al., 2019). We then harmonized species names across niche, range, and phylogeny datasets, retaining only species with matched binomials across all sources. Additionally, we compiled generation length estimates for all taxa (Mancini et al., 2025; Soria et al., 2021; Tobias et al., 2022), imputing a small proportion of missing values (1.8% birds; 0.8% mammals) using phylogenetically informed estimates derived from the same phylogenies with the *funspace* (v0.2.2) package (Carmona et al., 2024). We then downloaded and assigned threat statuses from the IUCN Red List to all species present in the dataset and generated two threat categories: Not Threatened (Least Concern [LC] and Data Deficient [DD]), and Threatened (Critically Endangered [CR], Endangered [EN], Vulnerable [VU] and Near Threatened [NT]) (Ali et al., 2023). Finally, we combined all observations of species’ niche limits, threat status, generation lengths, and range area estimates into a single dataset (Data S1).

### Realized niche limits and extinction risk

We obtained yearly mean thermal and aridity realized niche limits for 7,958 amphibian, 11,044 bird, 5,655 mammal, and 9,284 reptile species (Watson & Kerr, 2025a). For each species, we also extracted total geographic range area (km²) using expert-informed range maps (BirdLife International and Handbook of the Birds of the World, 2021; IUCN, 2022a). To account for shared evolutionary history in subsequent analyses, we obtained phylogenies for amphibians, birds, mammals, and reptiles from published sources (Jetz et al., 2012; Jetz & Pyron, 2018; Tonini et al., 2016; Upham et al., 2019). We then harmonized species names across niche, range, and phylogeny datasets, retaining only species with matched binomials across all sources. Additionally, we compiled generation length estimates for all taxa (Mancini et al., 2025; Soria et al., 2021; Tobias et al., 2022), imputing a small proportion of missing values (1.8% birds; 0.8% mammals) using phylogenetically informed estimates derived from the same phylogenies with the *funspace* (v0.2.2) package (Carmona et al., 2024). Wee then downloaded and assigned threat statuses from the IUCN Red List to all species present in the dataset and generated two threat categories: Not Threatened (Least Concern [LC] and Data Deficient [DD]), and Threatened (Critically Endangered [CR], Endangered [EN], Vulnerable [VU] and Near Threatened [NT]) (Ali et al., 2023). Finally, we combined all observations of species’ niche limits, threat status, generation lengths, and range area estimates into a single dataset.

### Species extinctions and phylogenetic diversity

We simulated extinctions across terrestrial vertebrate species (amphibians, birds, mammals, reptiles) for the next 100 years using the *iucnsim* (v0.0.0.9) R package (Andermann et al., 2021). This framework uses historical IUCN status transition rates and species-specific generation lengths to estimate annual probabilities of transitions between extinction risk categories. For each taxonomic group, we used the full IUCN-assessed species list as the reference set to estimate transition probabilities among threat categories. These probabilities determined simulated movement toward higher or lower extinction risk states over time independent of climate and land use variables. Importantly, these estimates reflect only IUCN-based transition dynamics informed by species-specific generation lengths.

From the harmonized niche limit dataset, we created a list of target species for generating extinction simulations by retaining all species with both IUCN status and niche information. We ran 100 independent simulations per taxonomic group, each projecting status transitions over 100 years, controlling for species generation length. We specifically simulated extinctions assuming no improvements to conservation efforts or increased threats to biodiversity. We extracted the list of species that reached extinction status in each simulation. Figure 6 displays the global distribution of proportional species lost averaged across all taxonomic groups and simulations. To generate a null expectation, we constructed matched random extinction scenarios. For each simulation, we randomly selected the same number of extinct species while preserving the proportion of Threatened and Not Threatened species. This produced paired extinction sets (IUCN-based and random) for each of the 100 simulations per taxonomic group. We then used the *caper* (v1.0.4) package (Orme et al., 2025) to quantify phylogenetic diversity (PD) as the sum total branch length of surviving species after pruning extinct species from the phylogenies (Faith, 1992). This generated 100 PD estimates for each extinction scenario (IUCN-based and random) and taxonomic class.

**Figure 6.**
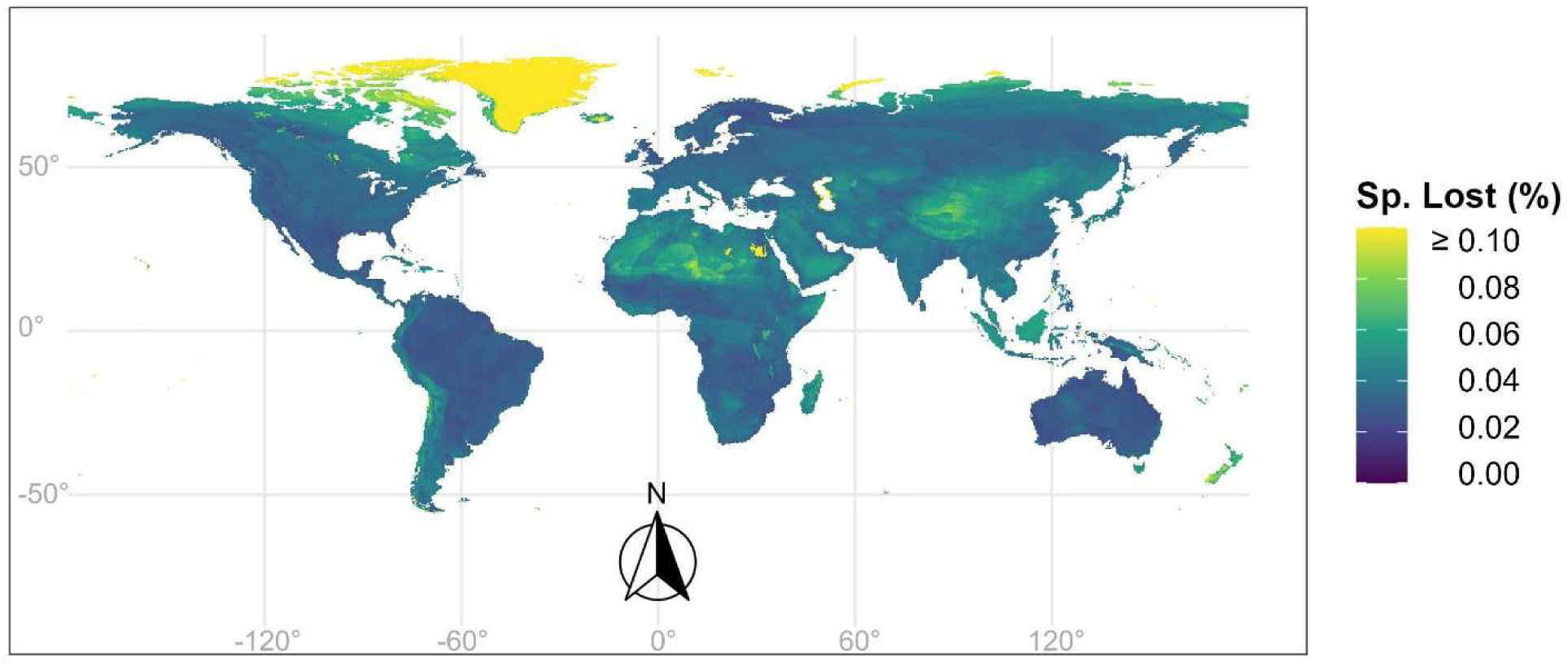
Distribuition of percent species lost under future IUCN simulated extinctions. The map displays the mean proporition of species simulated as extinct using the *iucnsim* package over the next 100 for terrestrail amphibians, birds, mammals, and reptiles at a gridded resoultion of 2.5 minutes (∼21 km2 at the equator). Map projection WGS84 (unprojected).

### Phylogenetic Signal

We tested whether realized thermal and aridity niche limits are phylogenetically conserved using Pagel’s λ implemented in the *geiger* package (v2.0.11) (Pennell et al., 2014). Pagel’s λ ranges from 0 (no phylogenetic signal) to 1 (trait variation consistent with Brownian motion evolution), although values greater than 1 can occur (Pearse et al., 2025). To evaluate alternative models of trait evolution, we also fitted Ornstein–Uhlenbeck (OU) models, which incorporate stabilizing selection toward adaptive optima (Butler & King, 2004). We compared models using AIC values to determine whether Brownian motion (reflected by Pagel’s λ) or OU processes better explained variation in realized niche limits across taxa.

### Statistical analyses

We tested whether realized niche limits predict current extinction risk using phylogenetic generalized linear models (PGLMs) implemented in the *phylolm* (v2.6.5) package (Tung Ho & Ané, 2014). This approach accounts for phylogenetic non-independence by incorporating class-specific phylogenetic covariance structures. For each taxonomic class we fit separate PGLMs, applying class specific phylogenetic distance matrices to the model error term. We model IUCN threat status (“Not Threatened” = 0, “Threatened” = 1) as a function of lower and upper thermal and aridity niche limits. Since our response variable was a binary categorical variable, we selected a binomial distribution with a logit link function. We also include log_10_-transformed range area as a covariate in our analyses since relationships between range size and extinction risk have previously been observed (Staude et al., 2020). All model predictor variables were z-transformed prior to analysis to allow for direct comparison between niche limit effects. All models were inspected and found to meet the assumptions of logistic regression analyses. When outliers were detected, we removed the observations from our dataset and generated new models using the filtered data. We found no influence of outlier observations on the direction of estimated trends or estimate significance across all models.

Next, we tested if realized niche limits differed between extinction scenarios (IUCN-based vs random) following a linear mixed effect model approach using the *glmmTMB* package (v1.1.11) (Brooks et al., 2017). We fit separate models for each niche limit variable which served as my response. The dataset used for these analyses included all species classed as extinct for all 100 IUCN and random extinction scenarios. Our predictor variables included extinction scenario, taxonomic class, and the interaction between them. To control for potential baseline differences in niche limits between extinction categories and avoid corollary analyses, we include threat status as a random intercept term. We also included simulation sample number as a random intercept term to control for repeat sampling of species across simulations. We again controlled for potential effects of range area by included log_10_ range area (km^2^) for each species as a covariate. For each niche limit model, we then extracted the estimated marginal means of the contrasts between IUCN and random simulations for each taxonomic class. All models were inspected and found to meet the assumptions of linear regression.

Finally, we used linear regression analysis to compare PD measurements between extinction scenarios. We used the previously generated PD estimates as my response variable, which was z-scaled separately for each taxonomic class prior to analysis. This approach controlled for potential differences in absolute branch lengths between taxonomic groups and had no impact on model results or interpretation. As in previous analyses, we then included taxonomic class, extinction scenario, and the interaction between them as my predictor variables. Finally, we again extracted the estimated marginal means for the contrast between IUCN and random extinction scenarios based on taxonomic group. The model was inspected and found to meet the assumptions of linear regression.

## Author details

**Matthew Nagy-Watson**

University of Ottawa, Ottawa, Canada

**Contribution**: Conceptualization, Data curation, Formal analysis, Validation, Investigation, Visualization, Methodology, Writing - original draft, Writing – review and editing

**Jeremy Kerr**

University of Ottawa, Ottawa, Canada

**Contribution**: Conceptualization, Methodology, Resources, Software, Supervision, Funding acquisition, Validation, Writing – review and editing

## Data availability

All data and scripts needed to evaluate the conclusions in the paper are available through figshare (10.6084/m9.figshare.32718738).

## References

Albright, T. P., Mutiibwa, D., Gerson, A. R., Smith, E. K., Talbot, W. A., O’Neill, J. J., McKechnie, A. E., & Wolf, B. O. (2017). Mapping evaporative water loss in desert passerines reveals an expanding threat of lethal dehydration. Proceedings of the National Academy of Sciences, 114(9), 2283–2288. 10.1073/pnas.1613625114

Ali, J. R., Blonder, B. W., Pigot, A. L., & Tobias, J. A. (2023). Bird extinctions threaten to cause disproportionate reductions of functional diversity and uniqueness. Functional Ecology, 37(1), 162–175. 10.1111/1365-2435.14201

Allan, E., Manning, P., Alt, F., Binkenstein, J., Blaser, S., Blüthgen, N., Böhm, S., Grassein, F., Hölzel, N., Klaus, V. H., Kleinebecker, T., Morris, E. K., Oelmann, Y., Prati, D., Renner, S. C., Rillig, M. C., Schaefer, M., Schloter, M., Schmitt, B., Schöning, I., Schrumpf, M., Solly, E., Sorkau, E., Steckel, J., Steffen-Dewenter, I., Stempfhuber, B., Tschapka, M., Weiner, C. N., Weisser, W. W., Werner, M., Westphal, C., Wilcke, W., & Fischer, M. (2015). Land use intensification alters ecosystem multifunctionality via loss of biodiversity and changes to functional composition. Ecology Letters, 18(8), 834–843. 10.1111/ele.12469

Andermann, T., Faurby, S., Cooke, R., Silvestro, D., & Antonelli, A. (2021). iucn_sim: A new program to simulate future extinctions based on IUCN threat status. Ecography, 44(2), 162–176. 10.1111/ecog.05110

Andreasson, F., Nord, A., & Nilsson, J.-Å. (2018). Experimentally increased nest temperature affects body temperature, growth and apparent survival in blue tit nestlings. Journal of Avian Biology, 49(2), Article jav-01620. 10.1111/jav.01620

Barnosky, A. D., Matzke, N., Tomiya, S., Wogan, G. O. U., Swartz, B., Quental, T. B., Marshall, C., McGuire, J. L., Lindsey, E. L., Maguire, K. C., Mersey, B., & Ferrer, E. A. (2011). Has the Earth’s sixth mass extinction already arrived? Nature, 471(7336), 51–57. 10.1038/nature09678

Bird, J. P., Martin, R., Akçakaya, H. R., Gilroy, J., Burfield, I. J., Garnett, S. T., Symes, A., Taylor, J., Şekercioğlu, Ç. H., & Butchart, S. H. M. (2020). Generation lengths of the world’s birds and their implications for extinction risk. Conservation Biology, 34(5), 1252–1261. 10.1111/cobi.13486

BirdLife International, & Handbook of the Birds of the World. (2021). Bird species distribution maps of the world (Version 2021.1). http://datazone.birdlife.org/species/requestdis

Boyles, J. G., Seebacher, F., Smit, B., & McKechnie, A. E. (2011). Adaptive thermoregulation in endotherms may alter responses to climate change. Integrative and Comparative Biology, 51(5), 676–690. 10.1093/icb/icr053

Brooks, M. E., Kristensen, K., van Benthem, K. J., Magnusson, A., Berg, C. W., Nielsen, A., Skaug, H. J., Mächler, M., & Bolker, B. M. (2017). glmmTMB balances speed and flexibility among packages for zero-inflated generalized linear mixed modeling. The R Journal, 9(2), 378–400. 10.32614/RJ-2017-066

Brooks, T. M., Mittermeier, R. A., Mittermeier, C. G., Da Fonseca, G. A. B., Rylands, A. B., Konstant, W. R., Flick, P., Pilgrim, J., Oldfield, S., Magin, G., & Hilton-Taylor, C. (2002). Habitat loss and extinction in the hotspots of biodiversity. Conservation Biology, 16(4), 909–923. 10.1046/j.1523-1739.2002.00530.x

Buckley, L. B., Ehrenberger, J. C., & Angilletta, M. J. (2015). Thermoregulatory behaviour limits local adaptation of thermal niches and confers sensitivity to climate change. Functional Ecology, 29(8), 1038–1047. 10.1111/1365-2435.12406

Butler, M. A., & King, A. A. (2004). Phylogenetic comparative analysis: A modeling approach for adaptive evolution. The American Naturalist, 164(6), 683–695. 10.1086/426002

Calosi, P., Bilton, D. T., & Spicer, J. I. (2007). Thermal tolerance, acclimatory capacity and vulnerability to global climate change. Biology Letters, 4(1), 99–102. 10.1098/rsbl.2007.0408

Carmona, C. P., Pavanetto, N., & Puglielli, G. (2024). funspace: An R package to build, analyse and plot functional trait spaces. Diversity and Distributions, 30(4), e13820. 10.1111/ddi.13820

Ceballos, G., Ehrlich, P. R., Barnosky, A. D., García, A., Pringle, R. M., & Palmer, T. M. (2015). Accelerated modern human-induced species losses: Entering the sixth mass extinction. Science Advances, 1(5), e1400253. 10.1126/sciadv.1400253

Chen, I.-C., Hill, J. K., Ohlemüller, R., Roy, D. B., & Thomas, C. D. (2011). Rapid range shifts of species associated with high levels of climate warming. Science, 333(6045), 1024–1026. 10.1126/science.1206432

Conenna, I., Santini, L., Rocha, R., Monadjem, A., Cabeza, M., & Russo, D. (2021). Global patterns of functional trait variation along aridity gradients in bats. Global Ecology and Biogeography, 30(5), 1014–1029. 10.1111/geb.13278

D’Agostino, E. R. R., Vivero, R., Romero, L., Bejarano, E., Hurlbert, A. H., Comeault, A. A., & Matute, D. R. (2022). Phylogenetic climatic niche conservatism in sandflies (Diptera: Phlebotominae) and their relatives. Evolution, 76(10), 2361–2374. 10.1111/evo.14580

De Vos, J. M., Joppa, L. N., Gittleman, J. L., Stephens, P. R., & Pimm, S. L. (2015). Estimating the normal background rate of species extinction. Conservation Biology, 29(2), 452–462. 10.1111/cobi.12380

Di Marco, M., Collen, B., Rondinini, C., & Mace, G. M. (2015). Historical drivers of extinction risk: Using past evidence to direct future monitoring. Proceedings of the Royal Society B: Biological Sciences, 282(1813), 20150928. 10.1098/rspb.2015.0928

du Plessis, K. L., Martin, R. O., Hockey, P. A. R., Cunningham, S. J., & Ridley, A. R. (2012). The costs of keeping cool in a warming world: Implications of high temperatures for foraging, thermoregulation and body condition of an arid-zone bird. Global Change Biology, 18(10), 3063–3070. 10.1111/j.1365-2486.2012.02778.x

Faith, D. P. (1992). Conservation evaluation and phylogenetic diversity. Biological Conservation, 61(1), 1–10. 10.1016/0006-3207(92)91201-3

Faith, D. P. (2008). Threatened species and the potential loss of phylogenetic diversity: Conservation scenarios based on estimated extinction probabilities and phylogenetic risk analysis. Conservation Biology, 22(6), 1461–1470. 10.1111/j.1523-1739.2008.01068.x

Findell, K. L., Berg, A., Gentine, P., Krasting, J. P., Lintner, B. R., Malyshev, S., Santanello, J. A., & Shevliakova, E. (2017). The impact of anthropogenic land use and land cover change on regional climate extremes. Nature Communications, 8(1), Article 989. 10.1038/s41467-017-01038-w

Foden, W. B., Butchart, S. H. M., Stuart, S. N., Vié, J.-C., Akçakaya, H. R., Angulo, A., DeVantier, L. M., Gutsche, A., Turak, E., Cao, L., Donner, S. D., Katariya, V., Bernard, R., Holland, R. A., Hughes, A. F., O’Hanlon, S. E., Garnett, S. T., Şekercioğlu, Ç. H., & Mace, G. M. (2013). Identifying the world’s most climate change vulnerable species: A systematic trait-based assessment of all birds, amphibians and corals. PLOS ONE, 8(6), e65427. 10.1371/journal.pone.0065427

Gardner, J. L., Peters, A., Kearney, M. R., Joseph, L., & Heinsohn, R. (2011). Declining body size: A third universal response to warming? Trends in Ecology & Evolution, 26(6), 285–291. 10.1016/j.tree.2011.03.005

Golodets, C., Sternberg, M., Kigel, J., Boeken, B., Henkin, Z., Seligman, N. G., & Ungar, E. D. (2015). Climate change scenarios of herbaceous production along an aridity gradient: Vulnerability increases with aridity. Oecologia, 177(4), 971–979. 10.1007/s00442-015-3234-5

Hantak, M. M., McLean, B. S., Li, D., & Guralnick, R. P. (2021). Mammalian body size is determined by interactions between climate, urbanization, and ecological traits. Communications Biology, 4(1), 961. 10.1038/s42003-021-02505-3

Hargreaves, A. L., Samis, K. E., & Eckert, C. G. (2014). Are species’ range limits simply niche limits writ large? A review of transplant experiments beyond the range. The American Naturalist, 183(2), 157–173. 10.1086/674525

Harris, R. M. B., Beaumont, L. J., Vance, T. R., Tozer, C. R., Remenyi, T. A., Perkins-Kirkpatrick, S. E., Mitchell, P. J., Nicotra, A. B., McGregor, S., Andrew, N. R., Letnic, M., Kearney, M. R., Wernberg, T., Hutley, L. B., Chambers, L. E., Fletcher, M. S., Keatley, M. R., Woodward, C. A., Williamson, G., Baumgartner, J. B., & Bowman, D. M. J. S. (2018). Biological responses to the press and pulse of climate trends and extreme events. Nature Climate Change, 8(7), 579–587. 10.1038/s41558-018-0187-9

Helaouët, P., & Beaugrand, G. (2009). Physiology, ecological niches and species distribution. Ecosystems, 12(8), 1235–1245. 10.1007/s10021-009-9261-5

Hogue, A. S., & Breon, K. (2022). The greatest threats to species. Conservation Science and Practice, 4(5), e12670. 10.1111/csp2.12670

Huang, J., Yu, H., Guan, X., Wang, G., & Guo, R. (2016). Accelerated dryland expansion under climate change. Nature Climate Change, 6(2), 166–171. 10.1038/nclimate2837

IUCN. (2022a). The IUCN Red List of Threatened Species. https://www.iucnredlist.org

IUCN. (2022b). Threats classification scheme (Version 3.3). The IUCN Red List of Threatened Species. https://www.iucnredlist.org/resources/threat-classification-scheme

Jetz, W., & Pyron, R. A. (2018). The interplay of past diversification and evolutionary isolation with present imperilment across the amphibian tree of life. Nature Ecology & Evolution, 2(5), 850–858. 10.1038/s41559-018-0515-5

Jetz, W., Thomas, G. H., Joy, J. B., Hartmann, K., & Mooers, A. O. (2012). The global diversity of birds in space and time. Nature, 491(7424), 444–448. 10.1038/nature11631

Kerr, J. T., Gordon, S. C. C., Chen, I.-C., Ednie, G., Foden, W., Newbold, T., Reynolds, A. R., Suggitt, A. J., Terblanche, J. S., & Watson, M. J. (2025). Effects of microclimate variation on insect persistence under global change. Nature Reviews Biodiversity, 1, 1–11. 10.1038/s44358-025-00067-4

Kingsolver, J. G., Diamond, S. E., & Buckley, L. B. (2013). Heat stress and the fitness consequences of climate change for terrestrial ectotherms. Functional Ecology, 27(6), 1415–1423. 10.1111/1365-2435.12145

Lee-Yaw, J. A., Kharouba, H. M., Bontrager, M., Mahony, C., Csergő, A. M., Noreen, A. M. E., Li, Q., Schuster, R., & Angert, A. L. (2016). A synthesis of transplant experiments and ecological niche models suggests that range limits are often niche limits. Ecology Letters, 19(6), 710–722. 10.1111/ele.12604

Li, X., Hu, W., Bleisch, W. V., Li, Q., Wang, H., Lu, W., Sun, J., Zhang, F., Ti, B., & Jiang, X. (2022). Functional diversity loss and change in nocturnal behavior of mammals under anthropogenic disturbance. Conservation Biology, 36(3), e13839. 10.1111/cobi.13839

Liu, X., Guo, R., Xu, X., Shi, Q., Li, X., Yu, H., Ren, Y., & Huang, J. (2023). Future increase in aridity drives abrupt biodiversity loss among terrestrial vertebrate species. Earth’s Future, 11(4), e2022EF003162. 10.1029/2022EF003162

Luedtke, J. A., Chanson, J., Neam, K., Hobin, L., Maciel, A. O., Catenazzi, A., Borzée, A., Hamidy, A., Aowphol, A., Jean, A., Sosa-Bartuano, Á., Fong G, A., de Silva, A., Fouquet, A., Angulo, A., Kidov, A. A., Muñoz Saravia, A., Diesmos, A. C., Tominaga, A., Rustamov, E. A., Iskandar, D. T., Bickford, D., Brown, R. M., Chan, K. O., Dutta, S. K., Ohler, A., Richards, S. J., Rowley, J. J. L., Van Dijk, P. P., Stuart, B. L., Wogan, G. O. U., & Stuart, S. N. (2023). Ongoing declines for the world’s amphibians in the face of emerging threats. Nature, 622(7982), 308–314. 10.1038/s41586-023-06578-4

Luo, D., Hu, Z., Dai, L., Hou, G., Di, K., Liang, M., Cao, R., & Zeng, X. (2023). An overall consistent increase of global aridity in 1970–2018. Journal of Geographical Sciences, 33(3), 449–463. 10.1007/s11442-023-2091-0

Mancini, G., Santini, L., Cazalis, V., Ficetola, G. F., Meiri, S., Roll, U., Silvestri, S., Pincheira-Donoso, D., & Di Marco, M. (2025). Generation length of the world’s amphibians and reptiles. Ecography, 2025(7), e07527. 10.1111/ecog.07527

Matthews, T. J., Triantis, K. A., Wayman, J. P., Martin, T. E., Hume, J. P., Cardoso, P., Faurby, S., Mendenhall, C. D., Dufour, P., Rigal, F., Cooke, R., Whittaker, R. J., Pigot, A. L., Thébaud, C., Jørgensen, M. W., Benavides, E., Soares, F. C., Ulrich, W., Kubota, Y., Moura, M., Ficetola, G. F., Borda-de-Água, L., Kissling, W. D., Hortal, J., Rosindell, J., Santos, A. M. C., & Sayol, F. (2024). The global loss of avian functional and phylogenetic diversity from anthropogenic extinctions. Science, 386(6717), 55–60. 10.1126/science.adk7898

Miller-Struttmann, N. E. (2024). Climate change predicted to exacerbate declines in bee populations. Nature, 628(8007), 270–271. 10.1038/d41586-024-00681-w

Newbold, T., Hudson, L. N., Hill, S. L. L., Contu, S., Lysenko, I., Senior, R. A., Börger, L., Bennett, D. J., Choimes, A., Collen, B., Day, J., De Palma, A., Díaz, S., Echeverria-Londoño, S., Edgar, M. J., Feldman, A., Garon, M., Harrison, M. L. K., Alhusseini, T., Ingram, D. J., Itescu, Y., Kattge, J., Kemp, V., Kirkpatrick, L., Kleyer, M., Correia, D. L. P., Martin, C. D., Meiri, S., Novosolov, M., Pan, Y., Phillips, H. R. P., Purves, D. W., Robinson, A., Simpson, J., Tuck, S. L., Weiher, E., White, H. J., Ewers, R. M., Mace, G. M., Scharlemann, J. P. W., & Purvis, A. (2015). Global effects of land use on local terrestrial biodiversity. Nature, 520(7545), 45–50. 10.1038/nature14324

Oliver, T. H., Isaac, N. J. B., August, T. A., Woodcock, B. A., Roy, D. B., & Bullock, J. M. (2015). Declining resilience of ecosystem functions under biodiversity loss. Nature Communications, 6, 10122. 10.1038/ncomms10122

Orme, D., Freckleton, R., Thomas, G., Petzoldt, T., Fritz, S., Isaac, N., & Pearse, W. (2025). caper: Comparative analyses of phylogenetics and evolution in R (R package version 1.0.4). https://CRAN.R-project.org/package=caper

Pacifici, M., Visconti, P., Butchart, S. H. M., Watson, J. E. M., Cassola, F. M., & Rondinini, C. (2017). Species’ traits influenced their response to recent climate change. Nature Climate Change, 7(3), 205–208. 10.1038/nclimate3223

Paquette, A., & Hargreaves, A. L. (2021). Biotic interactions are more often important at species’ warm versus cool range edges. Ecology Letters, 24(11), 2427–2438. 10.1111/ele.13864

Pearse, W. D., Davies, T. J., & Wolkovich, E. M. (2025). How to define, use, and interpret Pagel’s λ (lambda) in ecology and evolution. Global Ecology and Biogeography, 34(4), e70012. 10.1111/geb.70012

Pennell, M. W., Eastman, J. M., Slater, G. J., Brown, J. W., Uyeda, J. C., FitzJohn, R. G., Alfaro, M. E., & Harmon, L. J. (2014). geiger v2.0: An expanded suite of methods for fitting macroevolutionary models to phylogenetic trees. Bioinformatics, 30(15), 2216–2218. 10.1093/bioinformatics/btu181

Quintero, I., & Wiens, J. J. (2013). Rates of projected climate change dramatically exceed past rates of climatic niche evolution among vertebrate species. Ecology Letters, 16(8), 1095–1103. 10.1111/ele.12144

Ratnayake, H. U., Kearney, M. R., Govekar, P., Karoly, D., & Welbergen, J. A. (2019). Forecasting wildlife die-offs from extreme heat events. Animal Conservation, 22(4), 386–395. 10.1111/acv.12476

Raup, D. M. (1986). Biological extinction in earth history. Science, 231(4745), 1528–1533. 10.1126/science.11542058

Riddell, E. A., Iknayan, K. J., Hargrove, L., Tremor, S., Patton, J. L., Ramirez, R., Wolf, B. O., & Beissinger, S. R. (2021). Exposure to climate change drives stability or collapse of desert mammal and bird communities. Science, 371(6529), 633–638. 10.1126/science.abd4605

Rubalcaba, J. G., Gouveia, S. F., Villalobos, F., Olalla-Tárraga, M., & Sunday, J. (2023). Climate drives global functional trait variation in lizards. Nature Ecology & Evolution, 7(4), 524–534. 10.1038/s41559-023-02007-x

Sannolo, M., & Carretero, M. A. (2019). Dehydration constrains thermoregulation and space use in lizards. PLOS ONE, 14(7), e0220384. 10.1371/journal.pone.0220384

Scholes, R. J. (2016). Climate change and ecosystem services. Wiley Interdisciplinary Reviews: Climate Change, 7(4), 537–550. 10.1002/wcc.404

Sheridan, J. A., & Bickford, D. (2011). Shrinking body size as an ecological response to climate change. Nature Climate Change, 1(8), 401–406. 10.1038/nclimate1259

Simkins, A. T., Sutherland, W. J., Dicks, L. V., Hilton-Taylor, C., Grace, M. K., Butchart, S. H. M., Senior, R. A., & Petrovan, S. O. (2025). Past conservation efforts reveal which actions lead to positive outcomes for species. PLOS Biology, 23(3), e3003051. 10.1371/journal.pbio.3003051

Soria, C. D., Pacifici, M., Di Marco, M., Stephen, S. M., & Rondinini, C. (2021). COMBINE: A coalesced mammal database of intrinsic and extrinsic traits. Ecology, 102(6), e03344. 10.1002/ecy.3344

Soroye, P., Newbold, T., & Kerr, J. (2020). Climate change contributes to widespread declines among bumble bees across continents. Science, 367(6478), 685–688. 10.1126/science.aax8591

Spooner, F. E. B., Pearson, R. G., & Freeman, R. (2018). Rapid warming is associated with population decline among terrestrial birds and mammals globally. Global Change Biology, 24(10), 4521–4531. 10.1111/gcb.14361

Staude, I. R., Navarro, L. M., & Pereira, H. M. (2020). Range size predicts the risk of local extinction from habitat loss. Global Ecology and Biogeography, 29(1), 16–25. 10.1111/geb.13003

Sunday, J. M., Bates, A. E., & Dulvy, N. K. (2012). Thermal tolerance and the global redistribution of animals. Nature Climate Change, 2(9), 686–690. 10.1038/nclimate1539

Takamata, A. (2012). Modification of thermoregulatory response to heat stress by body fluid regulation. The Journal of Physical Fitness and Sports Medicine, 1(3), 479–489. 10.7600/jpfsm.1.479

Taylor, E. N., Diele-Viegas, L. M., Gangloff, E. J., Hall, J. M., Halpern, B., Massey, M. D., Rödder, D., Rollinson, N., Spears, S., Sun, B. J., & Telemeco, R. S. (2021). The thermal ecology and physiology of reptiles and amphibians: A user’s guide. Journal of Experimental Zoology Part A: Ecological and Integrative Physiology, 335(1), 13–44. 10.1002/jez.2396

Theodoridis, S., Fordham, D. A., Brown, S. C., Li, S., Rahbek, C., & Nogués-Bravo, D. (2020). Evolutionary history and past climate change shape the distribution of genetic diversity in terrestrial mammals. Nature Communications, 11(1), 2557. 10.1038/s41467-020-16449-5

Thuiller, W. (2004). Patterns and uncertainties of species’ range shifts under climate change. Global Change Biology, 10(12), 2020–2027. 10.1111/j.1365-2486.2004.00859.x

Tobias, J. A., Sheard, C., Pigot, A. L., Devenish, A. J. M., Yang, J., Sayol, F., Neate-Clegg, M. H. C., Alioravainen, N., Weeks, T. L., Barber, R. A., Walkden, P. A., MacGregor, H. E. A., Jones, S. E. I., Vincent, C., Phillips, A. G., Marples, N. M., Montaño-Centellas, F. A., Leandro-Silva, V., Claramunt, S., Cooney, C. R., Darski, B., Davidson, G. L., Freeman, B. G., Hughes, E. C., Ireland, L., Kelly, D. J., McGregor, R., Mitteroecker, P., O’Shea, W., Ottenburghs, J., Pyle, A., Sedláček, O., Street, S. E., Töpfer, T., Trisos, C. H., Weeks, B. C., Wilkinson, C. L., Wilson, A., & Schleuning, M. (2022). AVONET: Morphological, ecological and geographical data for all birds. Ecology Letters, 25(3), 581–597. 10.1111/ele.13898

Tonini, J. F. R., Beard, K. H., Ferreira, R. B., Jetz, W., & Pyron, R. A. (2016). Fully-sampled phylogenies of squamates reveal evolutionary patterns in threat status. Biological Conservation, 204, 23–31. 10.1016/j.biocon.2016.03.039

Tung Ho, L. S., & Ané, C. (2014). A linear-time algorithm for Gaussian and non-Gaussian trait evolution models. Systematic Biology, 63(3), 397–408. 10.1093/sysbio/syu005

Upham, N. S., Esselstyn, J. A., & Jetz, W. (2019). Inferring the mammal tree: Species-level sets of phylogenies for questions in ecology, evolution, and conservation. PLOS Biology, 17(12), e3000494. 10.1371/journal.pbio.3000494

Urban, M. C. (2024). Climate change extinctions. Science, 386(6726), 1123–1128. 10.1126/science.sdp4461

Van de Ven, T. M. F. N., Fuller, A., & Clutton-Brock, T. H. (2020). Effects of climate change on pup growth and survival in a cooperative mammal, the meerkat. Functional Ecology, 34(1), 194–202. 10.1111/1365-2435.13468

Vandermeer, J. (1972). Niche theory. Annual Review of Ecology and Systematics, 3, 107–132. 10.1146/annurev.es.03.110172.000543

Watson, M., & Kerr, J. (2025a). Global dataset for realized thermal and aridity niche limits for terrestrial vertebrates. Scientific Data, 12(1), 1829. 10.1038/s41597-025-06130-1

Watson, M. J., & Kerr, J. T. (2025b). Climate-driven body size changes in birds and mammals reveal environmental tolerance limits. Global Change Biology, 31(5), e70241. 10.1111/gcb.70241

Williams, J. J., Freeman, R., Spooner, F., & Newbold, T. (2022). Vertebrate population trends are influenced by interactions between land use, climatic position, habitat loss and climate change. Global Change Biology, 28(3), 797–815. 10.1111/gcb.15978

Williams, J. J., & Newbold, T. (2021). Vertebrate responses to human land use are influenced by their proximity to climatic tolerance limits. Diversity and Distributions, 27(7), 1308–1323. 10.1111/ddi.13282

Williams, N. F., McRae, L., Freeman, R., Capdevila, P., & Clements, C. F. (2021). Scaling the extinction vortex: Body size as a predictor of population dynamics close to extinction events. Ecology and Evolution, 11(11), 7069–7079. 10.1002/ece3.7555

Winkler, K., Fuchs, R., Rounsevell, M., & Herold, M. (2021). Global land use changes are four times greater than previously estimated. Nature Communications, 12(1), 2501. 10.1038/s41467-021-22702-2

Woodroffe, R., Groom, R., & McNutt, J. W. (2017). Hot dogs: High ambient temperatures impact reproductive success in a tropical carnivore. Journal of Animal Ecology, 86(6), 1329–1338. 10.1111/1365-2656.12719

Woo-Durand, C., Matte, J.-M., Cuddihy, G., McGourdji, C. L., Grant, J. W. A., & Venter, O. (2020). Increasing importance of climate change and other threats to at-risk species in Canada. Environmental Reviews, 28(4), 449–456. 10.1139/er-2020-0032

Wu, N. C., Bovo, R. P., Enriquez-Urzelai, U., Clusella-Trullas, S., Kearney, M. R., Navas, C. A., & Kong, J. D. (2024). Global exposure risk of frogs to increasing environmental dryness. Nature Climate Change, 14(12), 1314–1322. 10.1038/s41558-024-02167-z

Zaifman, J., Shan, D., Ay, A., & Jimenez, A. G. (2017). Shifts in bird migration timing in North American long-distance and short-distance migrants are associated with climate change. International Journal of Zoology, 2017, 6025646. 10.1155/2017/6025646

Zhang, C., Li, L., Guan, Y., Cai, D., Chen, H., Bian, X., & Guo, S. (2021). Impacts of vegetation properties and temperature characteristics on species richness patterns in drylands: Case study from Xinjiang. Ecological Indicators, 133, 108417. 10.1016/j.ecolind.2021.108417

Zylstra, E. R., Rubke, C. A., & Steidl, R. J. (2025). Compounding effects of drought on long-term demography of a threatened reptile. Journal of Arid Environments, 227, 105315. 10.1016/j.jaridenv.2024.105315

